# The Influence of Phosphorus and Transplanting Date on Onion (*Allium Cepa* L.) Quality Parameters and yield of bulb Under Irrigation at Adami Tulu Jedo Kombolcha Woreda, East showa zone, Oromia Region

**DOI:** 10.1101/2021.01.12.426332

**Authors:** Tilaye Anbes, Walelign Worku

## Abstract

A field experiment was showed to decide the influence of phosphorus level and transplanting date on quality parameters of onion at Adami Tulu Jedo Kombolcha Woreda, during 2017/18 season. The treatments consisted of four phosphorus levels (0, 20, 40 and 60 kg P ha^−1^) and three transplanting date (42, 49 and 56 days). The experiment was laid out in randomized complete block design with four replications. The result showed that phosphorus level and transplanting date significantly influenced bulb dry matter content, marketable bulb yield, medium size bulb yield. Among these parameters, marketable bulb yield was also significantly influenced by the interaction of phosphorus level and transplanting date. On the other hand, small size bulb yield, large size bulb yield, over size bulb yield and under size bulb yield were only influenced by the effect of phosphorus levels. In this study, fertilized 60 kg P ha^−1^ with transplanting at 56 days of transplanting date recorded the highest marketable bulb yield, but no significant difference was showed with that obtained at 40 kg P ha^−1^ with the same transplanting date. Treatment combinations of no P (control) and transplanting date at 42 days produced the lowest amounts of marketable bulb yield. The economic analysis revealed that the highest net benefit with low cost of production was obtained in response to the application of 40 kg P ha^−1^ and the transplanting age of 56 days. The marginal rate of return for this treatment was 11983% which is found to be economically feasible for producing bulb in the districts.

## Introduction

Onion (*Allium cepa* L.) belongs to the family Alliaceae genus Allium (Griffiths et al., 2002), and it is a cross-pollinated, an herbaceous, biennial crop and pungent odor. It is one of the most important cool season vegetable crops. It ranks seconds among all vegetables in economic importance after tomatoes in the world (Griffiths et al., 2002; Mallor et al., 2011)

Onion is grown from seed, transplants or sets for use as both green onions and dry onion (Decoteau, 2000). It is used in several ways as in fresh, frozen, canned, pickled, powdered, chopped and dehydrated forms. Onions are primarily consumed for their unique flavor to enhance the flavor of other foods (Randle and Ketter, 1998), in powder form as spice for seasoning in cooking (Bagali et al., 2012). In addition, onion is known for its anti-bacterial, anti-viral, anti-allergenic and anti-inflammatory potential (Saha, 2013; Yemane et al., 2013). The mature bulb contains some starch, appreciable quantities of sugars, some protein, and vitamins A, B and C (Jilani *et al*., 2010).

The average productivity of onion in Ethiopia is 10.1 t ha^−1^ (CSA, 2015). This is very low yield compared to the world average of 19.3 t ha^−1^ (FAOSTAT, 2013). The low yield level could be due to low soil fertility, salinity effect and inappropriate transplanting and cultural practice (MARC, 2004). Geremew et al. (2010) suggested the control of plant spacing is one of the cultural practices to control bulb size, shape and yield. In the study area, onion crop plays an important role in contributing to the household food security. In addition to the nutritional value, these crops generate employment opportunities for the poor households in the district area.

The effect of transplant date on yield is an issue often broached by growers of horticultural and agronomic crops in an effort to maximize production potential (Leskovau and Vavrina, 1999). Although production of Bombay Red onion variety is expanding, information on Optimum phosphorus application rate and exact transplanting date is flimsy. Efficient investigation on fertilization to improve the quality parameters are missing. Onion producers in the area use comprehensive recommendation of phosphorus fertilizer which was recommended at country level. On the other hand farmers transplant onion based on their own judgment on the size of seedlings which critically influence the productivity and quality of the bulb. Both late and early date transplanting of seedlings may have significant influence on survival and quality of bulb yield. In view of the existing problem, this study was proposed with the objective to decide the influence of transplanting date and phosphorus levels on quality and marketable bulb yield.

## Materials and Methods

### Description of study area

The study was conducted at Adami Tulu Jedo Kombolcha Woreda during the 2017/18 dry season under irrigated condition. The site is located 220 km south of Addis Ababa city and 34 km west of Bulbula town in the vicinity of Abidjata and Shalla lakes. It is situated between 7^0^ 65’ N latitude and 38^°^ 56’ E longitudes and at an altitude of 1500 meters above sea level in the agro ecology of dry plateau of the Eastern part of the Ethiopian rift valley system. High amount of rainfall is received in the month of July and August. While the mean annual rainfall is 810 mm, the annual mean minimum and maximum temperatures are 12 ^°C^ and 30 ^°C^, respectively (Agerie and Afework, 2013).

### Experimental Planting material

The plant material for this study was Bombay Red variety of onion. The variety is widely accepted by farmers for its early maturity and higher bulb yield. It was released by Melkasa Agricultural Research Center (MARC) in 1980. It is well adapted to areas of 700-2000 meter above sea level (EARO, 2004). It is one of the most commonly and widely used improved variety in the Rift Valley System of Ethiopia and particularly at Adami Tulu Gedo Kombolcha Woreda.

### Treatments and Experimental Design

The experiment was contained of 4 × 3 factorial combinations involving phosphorus level and transplanting date. four P level (0, 20, 40, and 60 kg P ha^−1^) and Three transplanting date (42, 49 and 56 days of transplanting date) were laid out in randomized complete block design (RCBD) with four replications. Each treatment arrangement was allocated randomly to the experimental units within a block. Double row planting was done on ridges of about 15 cm height adopting recommended spacing of 40 cm between water furrows, 20 cm between rows on the ridge and 10 cm between plants within the row. The unit plot size of the experiment was 2 m x 2.3 m (4.6 m^2^). The blocks were parted by a distance of 1.0 m whereas the space between each plot within a block was 50 cm. In each plot, there were 10 rows, and in each row there were 20 plants. Totally, there were 200 plants per plot. The outer two rows at both sides of the plot and two plants at both ends of the rows were considered as border plants. The plants in the six central rows were used as net plot area to decide yield per plot and other quality parameters.

### Agronomic Practices and Treatment Applications

#### Raising onion seedlings

Seedlings of Bombay Red onion variety were raised in a nursery at Alage ATVET College demonstration site on sunken beds with size of 1 m x 5 m. The seed of Bombay Red was gained from Melkasa Agricultural Research Center, Horticulture Division. After four sunken nursery beds were prepared, seeds were sown on December 15/2017. The seed was drilled in well pulverized sunken bed in rows 10 cm apart and lightly covered with soil in the required transplanting date. All important cultural practices such as application of fertilizer (Urea and NPS), watering, weed, diseases and insect pest control activities, respectively were carry out based on the recommendations made for the onion crop.

### Experimental field management

Before transplanting seedlings, the experimental field was ploughed and leveled by tractor; ridges and plots were prepared manually. Large clods were broken down in order to bring the land to a fine tilth, and then a total of 48 plots based on recommended size were prepared in which 12 plots were allocated in each of the four replications. Moreover, the vital numbers of ridges and rows were marked in each plot. The seedlings were grown in the nursery with careful management and strict follow up until seedlings reached to the required stage as per the treatments. A three to four days irrigation interval was maintained for the first four weeks. Thereafter, irrigation was applied at seven days interval until fifteen days remaining to harvest, when irrigation was stopped completely (EARO, 2004).

As per the recommendation made for the onion in the study area half 50 kg ha^−1^ dose of N was applied uniformly to all plots during transplanting. The remaining half 50 kg ha^−1^ dose of nitrogen rates was side-dressed 45 days after transplanting for all plots (EARO, 2004; SARC, 2008; Anisuzzaman *et al*., 2009). Phosphorus (TSP) was applied as a single application as per specified rates at the time of transplanting based on the treatments. For the control of disease (purple blotch) and insect pest (onion thrips) the fungicides, Mancozeb 80 WP (3 kg ha^−1^) plus Ridomil (3.5 kg ha^−1^) and the insecticide, Selecron 720 EC (0.5 l ha^−1^) were used, respectively. All other agronomic practices were applied uniformly for all the plots as per the recommendation made for the crop (EARO, 2004).

### Soil Sampling

Before planting soil sample was taken randomly in a diagonal fashion from the experimental field at the depth of 0-30 cm for determination of physical and chemical properties of the soil. Nine soil samples were collected using an auger from the whole experimental field and combined to form a composite sample in a bucket. From this mixture, a sample weighing one kg was packed in to a plastic bag. The soil samples were also analyzed for soil texture, total nitrogen, cation exchange capacity (CEC), exchangeable potassium, organic carbon and available phosphorous. All the analyses were made at Zeway soil and water analysis laboratory in Batu Town.

### Data Collection

Dry matter content, total soluble solid and pyruvate concentration were recorded from twelve sample bulbs randomly taken from six central middle rows of each experimental plot. However, all plants in each net plot were harvested to collect data for bulb yield.

### Economic Analysis

Economic analysis was conducted to evaluate the economic feasibility of the treatments. Partial budget, dominance and marginal analysis were used. The analysis was based on data collected from respective district office of trade and transport, cooperatives and from onion fields. At Adami Tulu Jedo Kombolcha Woreda, the cost of 100 kg phosphorus (TSP) was 1200 birr and onion price of 400 birr per 100 kg was used for the net benefit analysis. The economic analysis is a way of calculating the total costs that vary and the net benefits of each treatment (CIMMYT, 1988).

### Data Analysis

The collected data on quality parameters under study were subjected to analysis of variance (ANOVA) using the GLM procedures of SAS (Statistical Analysis System) version 9.2 computer software program (SAS Institute Inc, 2008). Significance of differences between means was stated using the least significance difference (LSD) test at P< 0.05 probability level.

## Results and Discussion

### Experimental Soil Properties

The soil had particle size distribution of 18% sand, 36% silt and 46% clay at the depth of 0 - 30 cm. The results revealed that the texture of the composite soil sample from the site was silty clay loam. The pH was slightly alkaline (pH=7.83). However, the pH of this soil is near optimum for onion crop production although it is not ideal. According to the rating of Walkley and Black (1954) and Dewis and Freitas (1975), the soil of the study area is medium in organic carbon (2.10%) as well as total nitrogen (0.17%), respectively. The cation exchange capacity (CEC) (28.62 meq/100g) of the experimental soil is also high according to the rating of Jackson (1975), and low in phosphorus (3.87 mg/kg) according to Olsen *et al*. (1954).

### Quality Parameters

#### Bulb dry matter content

The main effect of phosphorus (P < 0.001) and transplanting date (P < 0.05) significantly affected mean bulb dry matter content of onion plants, however, the interaction effect did not show significant differences (P > 0.05). This study indicated that, increasing the rate of phosphorus from nil to 20 kg P ha^−1^ not significantly increased the bulb dry matter content of onion. But, increasing the rate of phosphorus further from 20 to 40 kg ha^−1^ increased the dry matter content of onion plants. And also the mean dry matter content of plants did not show significant difference as further increase in phosphorus rate from 40 to 60 kg P ha^−1^. Thus, the mean dry matter content of onion treated with phosphorus at the rate of 40 kg ha^−1^ exceeded the bulb dry matter of onion plants treated with nil and 20 kg P ha^−1^ by about 15 and 13%, respectively.

Similar observations were reported by Woldetsadik (2003) who stated that on a clay soil in a sub-humid tropical environment of Ethiopia in shallot (*Allium ascalonicum*) crop, increased application of P slightly increased bulb dry matter content of onion. Similarly, Tibebu *et al*. (2014) reported that the rate of P application increased dry matter content of onion.

Significant differences were observed among the transplanting date levels, in such a way that the 56 days of transplanting date gave the highest onion bulb dry matter content, while the lowest bulb dry matter content was obtained under 42 days of transplanting date. However, no statically significant difference was observed between 49 and 56 days of transplanting date (Table 1). The dry matter content for different date of transplanting varied possibly due to variation of growth patterns and photosynthesis at growing phases. The results of the present study are in agreement with Latif *et al*. (2010), Sultana (2015) and Bhonde *et al*. (2001) who reported that dry matter content of onion bulb was significantly influenced by the date of transplanting. This might be due to the fact that the optimum date of transplanting planted had better growth, which resulted in higher production of dry matter content of bulb (Sultana, 2015). This result is also consistent with the findings of Muhammad *et al*. (2016) who reported that the seedling transplanted at 60 days have high dry matter percentage as compared to the seedling transplant in early stage and it might be attributed to the fact that as the bulb size decreased quantity of water content also decreased resulting in high percentage of dry matter.

**Table 1:** Main effect of transplanting date and phosphorus levels on mean Bulb dry matter content (%), Total soluble solid(%) and Pyruvate concentration.

| Treatments | Bulb dry matter content (%) | Total soluble solids (%) | Pyruvate ($\mu\text{mole ml}^{-1}$) |
| --- | --- | --- | --- |
| **P ($\text{kg ha}^{-1}$)** |  |  |  |
| 0 | 11.67b | 12.05 | 21.23 |
| 20 | 11.83b | 11.96 | 20.91 |
| 40 | 13.44a | 12.29 | 19.56 |
| 60 | 14.26a | 12.13 | 20.96 |
| **LSD(0.05)** | 1.25 | 0.37 | 0.53 |
| **Significance level** | ** | NS | NS |
| **Transplanting date (days)** |  |  |  |
| 42 | 11.78b | 12.01 | 19.41c |
| 49 | 12.69a | 12.13 | 19.82ab |
| 56 | 13.46a | 12.00 | 21.38a |
| **LSD(0.05)** | 0.89 | 0.27 | 1.44 |
| **Significance level** | * | NS | * |
| **CV (%)** | 9.53 | 6.05 | 12.09 |

Means followed by the same letters within a column are not significantly different at ( $P < 0.05$ ), LSD 0.05

### Pyruvate concentration

Pyruvate concentration of bulb was significantly (P < 0.05) affected by the main effects of transplanting date treatments, but not significantly affected by the phosphorus levels and interaction effects of the treatments. The pyruvic acid (pyruvate) concentration of bulbs was significantly increased through the days of 49 and 56 transplanting date over the 42 days. Transplanting of onion at the days of 49 and 56 days has increased the pyruvate concentration by about 10 and 11%, respectively, over the 42 days. Randle and Ketter (1998) reported that to a large extent, the concentrations of pyruvate in onions are determined by the genetics of the cultivar. However, the growing environment can greatly influence the pungency intensity for any given cultivar.

### Total soluble solid (%)

Phosphorus fertilization and transplanting date and their interaction did not significantly (p > 0.05) affect the formation of total soluble solid (%) of bulb (Table 1). This could be due to the minimal direct effect of fertilization in the formation of total soluble solid bulbs.

### Small-sized bulb yield (20-50g)

The analysis of variance of small bulb size distribution of onion showed that the main effect of phosphorus fertilizer application had a significant influence (P < 0.001). However, neither the main effect of transplanting date nor its interaction with phosphorus influenced this parameter of onion.

Increasing the phosphorus application rate significantly decreased the production of small sized bulb yield as presented in Table 2. Thus, the highest small sized bulb yield was obtained from onion plants grown at the rate of 0 kg P ha^−1^. In contrast, the lowest small sized bulb yield of onion was recorded in response to the application of higher phosphorus rate at 60 kg ha^−1^ and 40 kg ha^−1^. For instance, the small sized bulb yield obtained in response to the application of 0 kg P ha^−1^ exceeded small sized bulb yield of plants grown with 60 kg ha^−1^ application by 96% (Table 2). The increment in small size bulb yield of onion in response to the application of nil phosphorus rates may have resulted in reduction in below ground bulb size like bulb weight, bulb length and diameter due to less availability of nutrients.

**Table 2:** Main effect of phosphorus levels and transplanting date on mean small size bulb yield, medium size bulb yield, large size bulb yield, over size bulb yield and under sized bulb yield.

| Treatments | Small size<br>bulb yield(t<br>ha<sup>-1</sup>) | Medium size<br>bulb yield(t<br>ha<sup>-1</sup>) | Large size<br>bulb<br>yield(t ha<sup>-1</sup>) | Under<br>sized bulb<br>yield (t<br>ha<sup>-1</sup>) | over size<br>bulb yield<br>(t ha<sup>-1</sup>) |
| --- | --- | --- | --- | --- | --- |
| **P (kg ha<sup>-1</sup>)** |  |  |  |  |  |
| 0 | 5.93<sup>a</sup> | 11.2<sup>c</sup> | 5.02<sup>c</sup> | 3.1<sup>a</sup> | 1.75<sup>c</sup> |
| 20 | 4.26<sup>b</sup> | 21.37<sup>b</sup> | 12.93<sup>b</sup> | 1.80<sup>b</sup> | 4.52<sup>b</sup> |
| 40 | 3.16<sup>c</sup> | 25.10<sup>a</sup> | 23.36<sup>a</sup> | 1.71<sup>b</sup> | 6.15<sup>a</sup> |
| 60 | 3.03<sup>c</sup> | 26.12<sup>a</sup> | 24.02<sup>a</sup> | 1.42<sup>b</sup> | 6.96<sup>a</sup> |
| **LSD(0.05)** | 0.76 | 2.66 | 3.15 | 0.59 | 0.89 |
| **Significance level** | *** | *** | *** | *** | *** |
| **Transplanting date**<br>**(days)** |  |  |  |  |  |
| 42 | 4.17 | 18.15<sup>b</sup> | 14.97 | 2.13 | 4.64 |
| 49 | 4.39 | 21.51<sup>a</sup> | 16.76 | 1.92 | 4.78 |
| 56 | 3.72 | 23.16<sup>a</sup> | 17.25 | 1.97 | 5.11 |
| **LSD(0.05)** | 0.66 | 2.30 | 2.73 | 0.51 | 0.77 |
| **Significance level** | NS | *** | NS | NS | NS |
| **CV (%)** | 22.35 | 15.28 | 23.22 | 35.35 | 22.07 |

Means followed by the same letters within a column are not significantly different at (P < 0.05), LSD 0.05

**Table 3:** Person correlation analysis of onion bulb yield and quality parameters at Adami Tulu Jedo Kombolcha Woreda.

|  | BDMC | MBY | SSBY | MSBY | LSBY | OSBY | USBY |
| --- | --- | --- | --- | --- | --- | --- | --- |
| BDMC | 1 |  |  |  |  |  |  |
| MBY | 0.00000 | 1 |  |  |  |  |  |
| SSBY | 0.00000 | 0.00000 | 1 |  |  |  |  |
| MSBY | 0.36285 | 0.60995 | 0.00477 | 1 |  |  |  |
| LSBY | 0.39798 | 0.61938 | -0.13736 | 0.53232 | 1 |  |  |
| OSBY | -0.12250 | -0.73423 | 0.21817 | -0.44911 | -0.66781 | 1 |  |
| USBY | 0.29894 | 0.79102 | 0.01679 | 0.49883 | 0.66704 | -0.61630 | 1 |

BDMC = bulb dry matter content, MBY= marketable bulb yield, SSBY= small sized bulb yield, MSBY= medium sized bulb yield, LSBY= large sized bulb yield, OSBY= oversized bulb yield and USBY= under sized bulb yield.

### Medium-sized bulb yield (50-100g)

Phosphorus fertilizer rate and Transplanting date exhibited highly significant (P < 0.001) variation on medium bulb size yield of onion. However, it was not significantly influenced by the interaction effect of those treatments.

This study indicated that, the production of medium sized bulb yield of onion was significantly increased by increasing the phosphorus application rate though the upper two rates were statistically at par. Hence, higher medium sized bulb yields were achieved from onion plants grown with the application of 60 kg P ha^−1^ and 40 kg P ha^−1^. On the other hand, zero application of phosphorus produced the lowest medium sized bulb yield (Table 2). This result is in agreement to the findings of Brewster (1994) who reported that increased phosphorus levels used to improve bulb size and increased the number of marketable bulbs in onion.

Significant differences were observed among the transplanting date treatments in such a way that the 56 days of transplanting date gave the highest medium sized bulb yield; while the lowest medium sized bulb yield was obtained under 42 days of transplanting date (Table 2). The size from 49 and 56 days age were not statistically different. The results are similar to the finding of Deepika (2013) who reported that yield of medium bulbs increased at optimum transplanting seedling age.

### Large-sized bulb yield (100-160g)

The analysis of variance showed that the main effect of phosphorus rate (P < 0.001) was significantly affected large sized bulb yield of onion. However, neither the main effect of transplanting date nor its interaction with phosphorus influenced this parameter of onion.

Similar to medium sized bulb yield, large sized bulb yield increased significantly in response to the increased application of phosphorus rate (Table 2). The maximum large sized bulb yield was obtained in response to the application of 60 kg P ha^−1^, but no significant different was observed with application rate of 40 kg P ha^−1^. On the other hand, the minimum value was achieved from the nil phosphorus rates.

### Over-sized bulb yield (>160g)

The analysis of variance showed that the main effect of phosphorus rate (P < 0.001) was significantly affected large sized bulb yield of onion. However, transplanting date and its interaction with phosphorus not significantly affected.

Similar to large sized bulb yield, oversized bulb yield increased significantly in response to the increased application of phosphorus rate (Table 2). The maximum oversized bulb yield was obtained in response to the application of 60 kg P ha^−1^, but no significant different was observed with application rate of 40 kg P ha^−1^. On the other hand, the minimum value was achieved from the control treatment.

### Under size bulb yield (< 20 g)

Results from the analysis of variance revealed that the phosphorus fertilizer rate was found to be significant (P < 0.01) on under sized bulb yield. However, the transplanting date and its interaction effect with phosphorus did not significantly affect under sized bulb yield of onion.

The maximum under sized bulb yield was recorded when onion plants received no P fertilizer. Conversely, the minimum under-sized bulb yield was recorded at the rates of 20, 40 and 60 kg P ha^−1^ (Table 2). Phosphorus application decreased under sized bulb yield though effects were similar among the three rates. The increase in the yield of the under-sized bulbs under unfertilized plots might be related with lack of P, which may have reduced vegetative growth like leaf number and length and bulb size by decreasing synthesis and partitioning of photosynthetic to the bulbs (Ghaffor *et al*., 2003). Phosphorus is important for root development and when unavailable plant growth is usually reduced. Its deficiency in onions reduces root and growth of leaf, size of bulb, yield and delay maturation (Abdissa *et al*., 2011).

Means followed by the same letters within a column are not significantly different at (P < 0.05), LSD 0.05

### Marketable bulb yield

Marketable bulb yield of onion was significantly affected (P < 0.001) by phosphorus levels and transplanting date. Similarly, significant interaction effect of transplanting date and phosphorus was perceived on the marketable bulb yield of onion (P<0.05).

Under 42 days of transplanting date, marketable bulb yield enhanced by about 100% at 42 kg ha^−1^ phosphorus compared to lowest yield recorded from unfertilized plots (figure 1). Further increase of phosphorus to 60 kg ha^−1^ did not significantly show variation, rather it showed a drop by about 30% and leveled off with yields from control plots and those fertilized at 20 kg ha^−1^ phosphorus. At 49 days of transplanting date, the maximum marketable bulb yield was produced at 40 kg ha^−1^ P level while the last marketable bulb yield was attained from control treatments. Significant differences were noted among yields at 20, 40 and 60 kg ha^−1^ P rate under the 49 days age. At 56 days of transplanting date, the control plot had notably reduced yield of marketable bulb as compared to the three phosphorus levels. The highest marketable bulb yield was recorded at 56 days of transplanting date combined with 60 kg ha^−1^ phosphorus rate; though statically at par to that obtained from 40 kg P ha^−1^ under similar transplanting date (figure 1).

**Figure 1.**
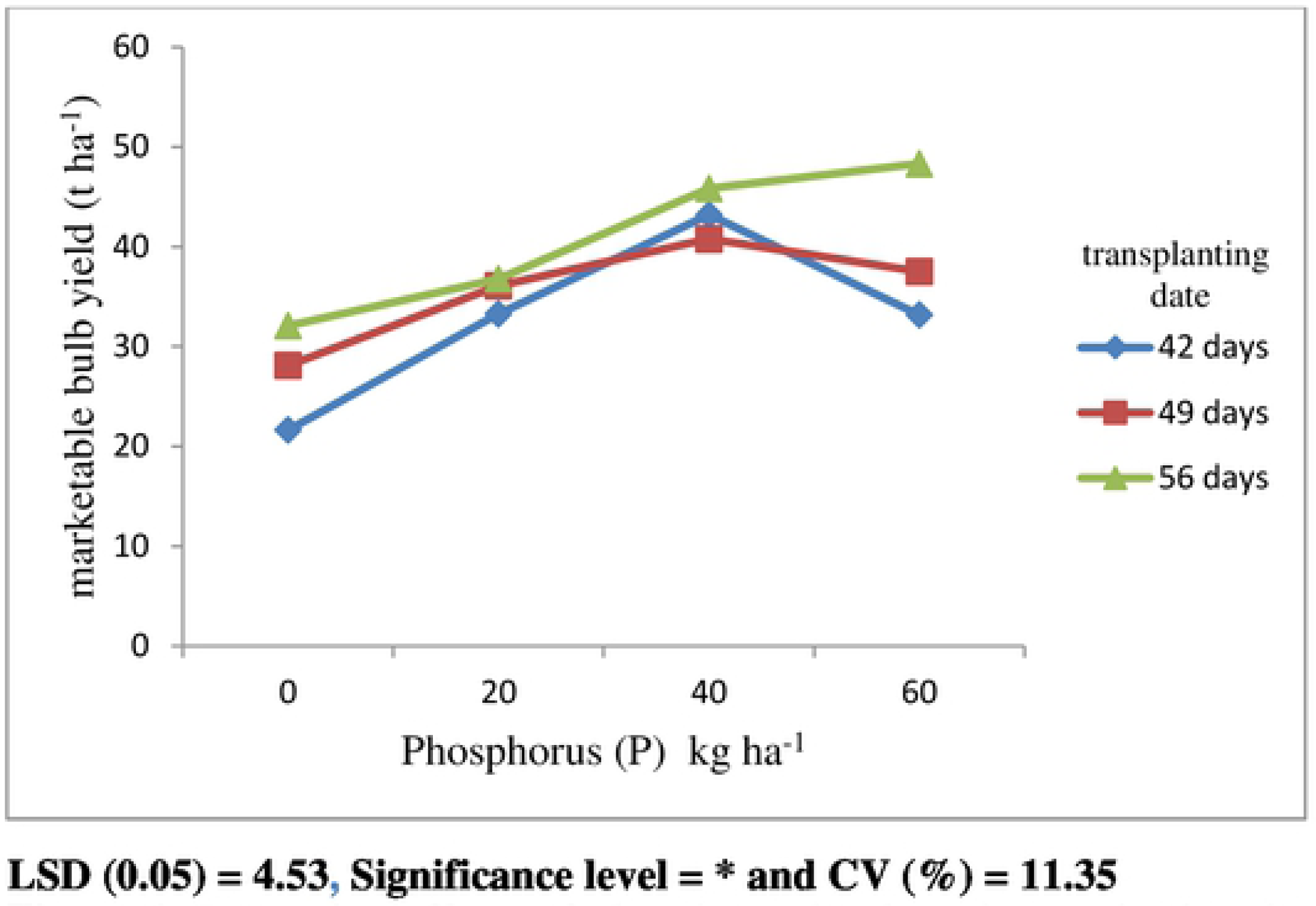
Interaction effects of phosphorus levels and transplanting date on marketable bulb yield (t ha^−1^) of onion grown at Adami Tutu Gedo Kombolcha Woreda.

From the current result it can be presumed that high transplanting date and higher phosphorus rate help to increase the vegetative growth of the plant which has developed assimilate availability for storage and led to an increased average bulb weight that gave an advantage to increase the marketable bulb yield.

### Economics Analysis

The economic analysis showed that the highest net benefit of Birr 146900 was recorded from the combination of 60 kg P ha^−1^ and 56 days of transplanting date with marginal rate of 3217%. This was followed by net benefit of Birr 139180 from the phosphorus rate of 40 kg P ha^−1^ and 56 days of transplanting date with the marginal rate of return of 11983%. This means that for every Birr 1.00 invested in 40 kg P ha^−1^ and 56 days of transplanting date, producers can expect to recover the Birr 1.00 and obtain an additional 119.83Birr. Whereas, the lowest net benefit (Birr 62480 ha^−1^) was recorded from control treatments (0 kg P ha^−1^) combined with 42 days of transplanting date (Table 4).

**Table 4:** Economic analysis for phosphorus and transplanting date experiments of onion at Adami Tulu Jedo Kombolcha Woreda.

| **Transplanting date with P (kg ha<sup>-1</sup>)** | **Average yield (t ha<sup>-1</sup>)** | **Adjusted yield (t ha<sup>-1</sup>)** | **Gross benefit (Birr ha<sup>-1</sup>)** | **Total cost that vary(Birr ha<sup>-1</sup>)** | **Net benefit (Birr ha<sup>-1</sup>)** |
| --- | --- | --- | --- | --- | --- |
| 42*20 | 31.15 | 24.92 | 99680 | 540 | 99140 d |
| 49*20 | 34.07 | 27.26 | 109040 | 540 | 108500 d |
| 56*20 | 34.68 | 27.74 | 110960 | 540 | 110420 |
| 42*40 | 41.12 | 32.89 | 131560 | 780 | 130780 d |
| 49*40 | 38.69 | 30.95 | 123800 | 780 | 123020 d |
| 56*40 | 43.74 | 34.99 | 139960 | 780 | 139180 |
| 42*60 | 31.07 | 24.86 | 99440 | 1020 | 98420 d |
| 49*60 | 35.41 | 28.33 | 113320 | 1020 | 112300 d |
| 56*60 | 46.23 | 36.98 | 147920 | 1020 | 146900 |
| 42*0 | 19.53 | 15.62 | 62480 |  | 62480 d |
| 49*0 | 26.08 | 22.94 | 91760 |  | 91760 d |
| 56*0 | 30.07 | 24.06 | 96240 |  | 96240 d |

Note: d= dominance

The minimum acceptable marginal rate of return (MRR %) should be between 50% and 100% (CIMMYT (1988)). Thus, the recent investigation indicated that marginal rate of return is higher than 100% (Table 5). Hence, the most economically attractive yield of the onion crop in the district area was that the combinations of 40 kg P ha^−1^ application and 56 days of transplanting date with low cost of production and higher benefits. The yield from the combination of 60 kg P ha^−1^with 56 days old seedlings still meets the 100% marginal rate of return threshold value. However, this experimental treatment will not be a viable alternative because the bulb yield from this combination was statistically at similar with that attained from 40 kg P ha^−1^and 56 days old age.

**Table 5:** Marginal analysis, phosphorus and transplanting date experiment in Adami Tulu Jedo Kombolcha Woreda.

| Transplanting date with P (kg ha<sup>-1</sup>) | Total cost that vary (Birr ha<sup>-1</sup>) | Marginal cost (Birr ha<sup>-1</sup>) | Net benefit (Birr ha<sup>-1</sup>) | Marginal net benefit (Birr ha<sup>-1</sup>) | Marginal rate of return (%) |
| --- | --- | --- | --- | --- | --- |
| 56*20 | 540 | - | 110420 | - | - |
| 56*40 | 780 | 240 | 139180 | 28760 | 11983 |
| 56*60 | 1020 | 240 | 146900 | 7720 | 3217 |

## Summary and Conclusions

The analysis of variance showed that medium size bulb yield, marketable bulb yield and bulb dry matter content were significantly influenced by the main effect of phosphorus rates and transplanting date. However, small size bulb yield, large size bulb yield, oversize bulb yield and under size bulb yield were significantly affected only by the main effects of different rates of phosphorus fertilizer. P fertilizer level and Transplanting date as well as their interaction did not significantly affect the formation of total soluble solid of bulb. From this study, significantly higher bulb dry matter content, medium size bulb yield were obtained at the transplanting date of 56 days and 40 kg P ha^−1^phosphorus rate. However, large sized bulb yield and oversized bulb yield were recorded in the treatments of 56 days of transplanting date and 60 kg ha^−1^ phosphorus rate. However, the result of this study showed that at 40 and 60 kg P ha^−1^ rate there was no significant variation in each parameter.

Therefore, the study revealed that, highest yield of marketable of bombay red onion variety was produced at treatment combination of 56 days of 60 kg ha^−1^ P rate with transplanting date, but no significant difference was observed in these parameters at 40 kg P ha^−1^ combinations with same transplanting date. However, the combination of 56 days of transplanting date fertilized with 40 kg ha^−1^ P rate also gave statistically comparable yield to the highest value. Therefore, from the present study it can be concluded that, the most economically attractive yield of the onion crop in the district area was obtained by the combinations of 40 kg P ha^−1^ applications and 56 days of transplanting date with low cost of production and higher benefits, however, to make reliable and acceptable recommendation it is better to repeat this experiment across locations and over seasons.

## Acknowledgement

The authors greatly would like thank you, all of you who are participated this piece of research work from the beginning to the end

## Conflict of Interests

The authors have not declared any conflicts of interests.

